# Structural abnormalities in thalamo-prefrontal tracks revealed by high angular resolution diffusion imaging predict working memory scores in concussed children

**DOI:** 10.1101/624445

**Authors:** Guido I. Guberman, Jean-Christophe Houde, Alain Ptito, Isabelle Gagnon, Maxime Descoteaux

**Affiliations:** Department of Neurology and Neurosurgery, Faculty of Medicine, Montreal Neurological Institute, McGill University, Quebec, Canada; Department of Computer Science, Sherbrooke University, Sherbrooke, Quebec, Canada; Department of Pediatrics, Faculty of Medicine, Montreal Children’s Hospital, McGill University, Quebec, Canada

**Keywords:** Concussion, mild TBI, Diffusion-Weighted Imaging, Constrained Spherical Deconvolution

## Abstract

Because of their massive prevalence, wide-ranging sequelae, and insidious nature, concussions are a potentially devastating neurological condition, especially in children. Shearing forces transmitted across the brain during concussions often result in white matter damage. The neuropathological impact of concussions has been discerned from animal studies and includes inflammation, demyelination, and axonal loss. These pathologies can overlap during the subacute stage of recovery. However, due to the challenges of accurately modelling complex white matter structure, these neuropathologies have not yet been differentiated in children *in vivo*. In the present study, we leveraged recent advances in diffusion imaging modelling, tractography, and tractometry to better understand the neuropathology underlying working memory problems in concussion. Studying a sample of 16 concussed and 46 healthy youths, we used novel tractography methods to isolate 11 working memory tracks. Along these tracks, we measured fractional anisotropy, diffusivities, track volume, apparent fiber density, and free water fraction. In three tracks connecting the right thalamus to the right dorsolateral prefrontal cortex (DLPFC), we found microstructural differences suggestive of myelin alterations. In another track connecting the left anterior-cingulate cortex with the left DLPFC, we found microstructural changes suggestive of axonal loss. Structural differences and tractography reconstructions were reproduced using test-retest analyses. White matter structure in the three thalamo-prefrontal tracks, but not the cingulo-prefrontal track, appeared to play a key role in working memory function. The present results improve understanding of working memory neuropathology in concussions, which constitutes an important first step towards developing neuropathologically-informed biomarkers of concussion in children.

## Introduction

Traumatic brain injury (TBI) is extremely common, affecting up to 600 per 100,000 individuals (Cassidy et al., 2004). The highest incidence rate occurs in early childhood and late adolescence (Faul, Xu, Wald, & Coronado, 2010). Its effect is wide-ranging, negatively impacting memory, mood, cognition, behaviour, and brain aging (Carroll et al., 2004; Dean, O’Neill, & Sterr, 2012; Kristman et al., 2014; Landre, Poppe, Davis, Schmaus, & Hobbs, 2006; McKee et al., 2009; Orlovska et al., 2014; Spinos et al., 2010). Mild forms of TBI (mTBIs or concussions, used here interchangeably), are the most common (Peeters et al., 2015) and are generally undetectable by conventional imaging such as Computerized Tomography (Lewine, Davis, Sloan, Kodituwakku, & Orrison, 1999; Shenton et al., 2012; Sundman, Doraiswamy, & Morey, 2015; Yuh et al., 2013) to the extent that cases of mTBI can often go undiagnosed (Meehan, Mannix, O’Brien, & Collins, 2013; Yuh et al., 2013). In cases of subtle or subclinical brain injuries, youths could be prematurely cleared to return to activities of daily life in the presence of incomplete cerebral recovery (Churchill et al., 2017) with potentially devastating consequences. Further, far fewer biomarker studies exist in children than in adults (Mayer et al., 2018). Neuroimaging biomarkers for pediatric concussions are still needed.

During a concussive event, the brain’s white-matter tends to be heavily affected (Bigler & Maxwell, 2012; Hayes, Bigler, & Verfaellie, 2016). Diffusion-tensor imaging (DTI) can provide different measures that describe water diffusion such as Fractional Anisotropy (FA), Axial Diffusivity (AD), Mean Diffusivity (MD), Radial Diffusivity (RD). These measurements are used to infer properties of white matter structure. A review of 100 DTI studies of TBI found that the vast majority of articles reported decreases in FA in injured individuals (Hulkower, Poliak, Rosenbaum, Zimmerman, & Lipton, 2013). Further, in a longitudinal study of 27 concussed varsity athletes, Churchill et al. (2017) found acute decreases in FA which remained present when patients were cleared to return to play. Hence, on long time scales, FA measurements can reliably detect abnormalities, even in the absence of symptoms.

Work from animal models has shown that shortly after impact, affected brain areas can display an edematous inflammatory response, accompanied by blood flow alterations, as well as metabolic abnormalities (Bigler & Maxwell, 2012). On longer time scales, as a result of these processes or in parallel, chronic neuroinflammation and edema, axonal demyelination, and/or axonal death can occur (Bigler & Maxwell, 2012; Pasternak, Kubicki, & Shenton, 2016). A DTI study using a mouse model of traumatic axonal injury showed that the presence of edema reduces FA (Mac Donald, Dikranian, Bayly, Holtzman, & Brody, 2007), which is known to result from an increase in free water (Pasternak et al., 2016). By decreasing diffusion hindrance, demyelination decreases FA (Budde, Janes, Gold, Turtzo, & Frank, 2011), and so does loss of axons (Harsan et al., 2006; S. W. Sun et al., 2006). However, in voxels with crossing fibers, loss of axons in one fiber population can paradoxically lead to an increase in FA (Douaud et al., 2011; Pierpaoli et al., 2001). During scarification, coherent astrocyte reactivity can also increase FA (Budde et al., 2011). Given its low specificity (Figure 1), FA cannot disambiguate between these different pathologies. As a result, individuals undergoing recovery at different rates can give rise to conflicting patterns of FA change across groups of concussed patients. A recent review focused on patients in sub-acute stages of recovery (less than 90 days) found, contrary to Hulkower et al. (2013), much more mixed results regarding the direction of FA change (Dodd, Epstein, Ling, & Mayer, 2014). Disambiguating the different forms of neuropathology present in patients at the sub-acute phase is therefore still needed given that each may have different clinical outcomes and require distinct treatment approaches.

**Figure 1.**
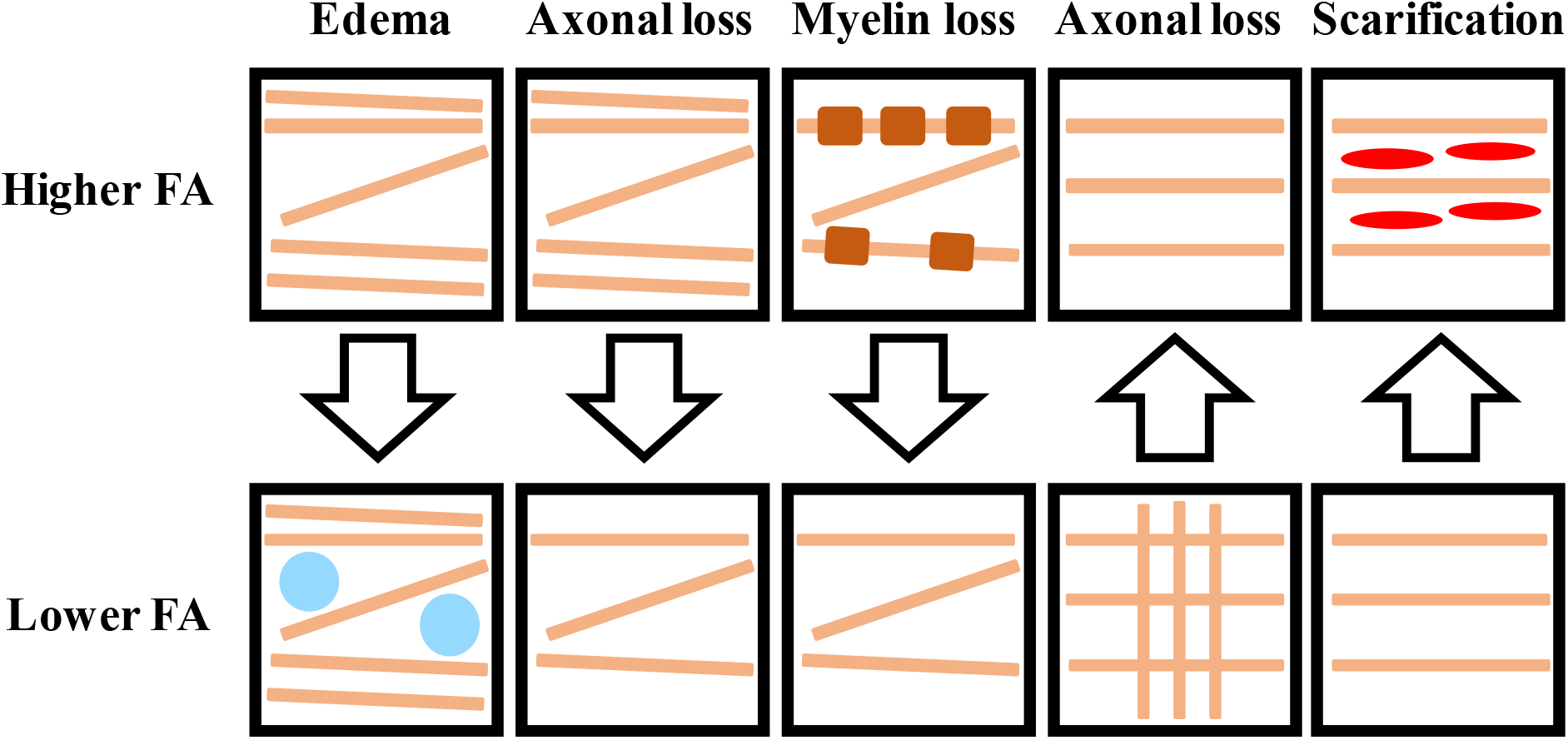
Illustration of the relative degree of FA in the context of different neuropathological processes. Arrows denote the direction of change as a result of injury.

Using other tensor-based measures has been proposed to complement FA and provide more information about mTBI neuropathology (Bigler & Maxwell, 2012; Budde et al., 2011). For instance, demyelination has been shown to increase RD and MD (Budde et al., 2011), whereas with loss of axons, a decrease in AD can be observed (Harsan et al., 2006; S. W. Sun et al., 2006). Despite these findings, interpretations of white matter microstructure from tensor-based measurements are limited (Jones, Knosche, & Turner, 2013), given that DTI assumes a single predominant fiber orientation per voxel (Tournier et al., 2008). Recent work suggests that this assumption is inaccurate across the majority of white matter voxels (Jeurissen, Leemans, Tournier, Jones, & Sijbers, 2013).

In the present study, we used advances in High Angular Resolution Diffusion Imaging (HARDI) local modelling, tractography, and tractometry to improve understanding of the neuropathology involved in concussion by using measures that, individually or in combination, provide information that is more specifically related to particular neuropathological processes. Given the wide scope of this question, we decided to focus on working memory deficits, a well-studied post-concussive abnormality. Previous work has repeatedly shown that concussed children and adults performing working memory tests display altered brain activity in the dorsolateral prefrontal cortex (DLPFC) as well as in other areas of the brain, irrespective of test performance (Chen, Johnston, Collie, McCrory, & Ptito, 2007; Chen et al., 2004; Jantzen, Anderson, Steinberg, & Kelso, 2004; Keightley et al., 2014; McAllister et al., 1999; McAllister et al., 2001; Pardini et al., 2010; Westfall et al., 2015). The areas of the brain related to working memory are well known, and focusing on this phenomenon allowed us to hone in on a small subset of 11 reliable tracks. Microstructural differences at the whole-track and segment-level were found for three right thalamo-prefrontal and one left cingulo-prefrontal tracks, suggestive of different, concurrent neuropathological processes. These differences were replicated in test-retest analyses using a second set of scans available for most participants. White matter structure in the thalamo-prefrontal tracks appeared to play a key role in working memory deficits in concussion.

## Materials and Methods

### Sample

Participants included 20 youths (11 females, 9 males; mean age=14.36 years, SD=2.65) who had previously sustained a concussion ranging from 9 to 96 days at the time of testing. Data from a group of 51 (20 females, 31 males, mean age=15.09, SD=1.89) healthy controls was obtained from previous imaging studies at the Montreal Neurological Institute. Participants were included in the study if they had received a diagnosis of mTBI, as per the World Health Organization task force definition (Cassidy et al., 2004) by a physician at the Montreal Children’s Hospital Concussion Clinic (Montreal, Quebec, Canada). All participants were screened prior to testing for a history of concussion, as well as to rule out other neurological conditions, including learning disabilities and ADHD. Participants were also excluded if they reported vestibular or musculoskeletal problems (other than upper extremity injuries), and if they had any contraindications for MRI, such as claustrophobia or metallic implants. Further information on recruitment and operational criteria for the identification of concussion are outlined in Keightley et al. (2014). No participants had abnormalities on standard imaging. Every participant provided informed written consent as approved by the ethics committee at McGill University, Montreal Neurological Institute.

### Neuropsychological testing

All participants completed a verbal and visual working memory test which consisted of an adapted version of the externally ordered working memory task, described elsewhere (Petrides, Frey, & Chen, 2001). This task has been validated in studies of patients and monkeys with lateral prefrontal lesions (Petrides, Alivisatos, Meyer, & Evans, 1993; Stern et al., 2000) and has been used in both concussed adults (Chen et al., 2004) and children to demonstrate reduced DLPFC activation, including in a subset of the present sample (Keightley et al., 2014). All participants also filled out a 21-item adapted version of the post-concussion symptom scale – revised (Lovell & Collins, 1998).

### Imaging Acquisition

All HARDI scans were acquired at the Montreal Neurological Institute using a 3.0T Siemens MAGNETOM Trio A Tim System with a 32-channel head coil. Each session started with acquisition of a high-resolution T1-weighted 3D anatomical image using 3D Magnetization Prepared Rapid Gradient Echo (MP-RAGE) sequence (TR = 23ms, TE = 2.98ms, Slice Thickness = 1mm, Image Matrix = 256 x 256, Flip Angle = 9 degrees, FOV = 256mm, interleaved excitation). DWI data were acquired with the following parameters: TR = 8300ms, echo time: 88ms, slice thickness = 2mm, FoV = 256mm, matrix = 128 x 128, number of slices=64, interleave excitation. Diffusion weighting was performed along 99 non-collinear directions using a b-value of 1000 s/mm^2^. Ten b=0s/mm^2^ images were acquired as reference. Out of 71 subjects, 67 underwent 2 DWI scans. All main analyses were performed on run 1 data. Run 2 data was used for replication purposes.

### Processing

A rigorous quality assessment procedure (see Supplementary Material) was performed, and images that passed were processed. Except where otherwise indicated, all steps in the processing pipeline were performed using the Diffusion Imaging for Python (DIPY) library and scripts built on it (Garyfallidis et al., 2014) (https://github.com/scilus/scilpy/). Images were first brain extracted using FSL’s *BET* command (Smith, 2002; Woolrich et al., 2009) and then denoised using Non-Local Spatial and Angular Matching, a novel tool that can be applied to any existing data and which improves the visual quality of data and results in more accurate and reproducible tractography results (St-Jean, Coupe, & Descoteaux, 2016). Images were then corrected for magnetic field inhomogeneity using Advanced Normalization Tools (ANTs) (Avants et al., 2011) and then eddy current and motion-corrected using FSL’s *eddy* command (Andersson & Sotiropoulos, 2016). Lastly, DWI images were upsampled to 1×1×1 space using trilinear interpolation (Girard, Whittingstall, Deriche, & Descoteaux, 2014; Tournier, Calamante, & Connelly, 2012), a procedure that, when performed before tractography, increases anatomical contrast, reduces partial volume effects and yields more accurate tract reconstructions (Dyrby et al., 2014).

### Local modelling

Free-water modelling was performed using the Accelerated Microstructure Imaging via Convex Optimization (AMICO) (Daducci et al., 2015) to calculate free-water fraction (FW), which is thought to be related to neuroinflammation (Pasternak et al., 2016; Pasternak, Shenton, & Westin, 2012). Processed DWI images were then corrected for FW, and water diffusion was modelled on FW-corrected DWI images at each voxel using a tensor. Outliers were then removed from FW-corrected AD, RD, and MD maps. Recent work suggests that removing outliers increases the robustness of tensor-based measurements (Jones et al., 2013). Then, fiber orientation distribution functions (fODFs) were computed using Constrained Spherical Deconvolution (CSD) (Descoteaux, Deriche, Knosche, & Anwander, 2009; Tournier, Calamante, & Connelly, 2007). Since fiber response functions (FRF) did not differ between groups, CSD was performed using an FRF averaged across all participants (Mito et al., 2018; Pierpaoli & Basser, 1996).

### Tractography

Probabilistic tractography was performed in upsampled DWI space using Particle Filtering Tractography (PFT) with anatomical priors. This method consists in optimizing tractography parameters by informing stopping criteria using partial volume estimation maps computed from high-resolution T1-weighted images which were transformed to upsampled DWI space using ANTs symmetric normalization (SyN) algorithm (Avants, Epstein, Grossman, & Gee, 2008; Avants et al., 2011). These maps provide an anatomically-informed, probabilistic stopping criterion, which, combined with a particle filtering method, can achieve tractography results that are less biased by length, shape, size, and position (Girard et al., 2014). The anatomical constraints used in this algorithm are: 1) the initiation of streamlines from the gray/white-matter interface; and 2) the obligatory termination of streamlines in gray matter. Gray/white-matter interface maps were created using FSL’s FAST (Y. Zhang, Brady, & Smith, 2001) in the transformed T1-weighted images. Seeding along the interface maps was performed using 24 seeds per voxel, which yielded an average of 2.3 million streamlines (SD=7.58*10^5^). Tractograms were cleaned by removing loops (defined as streamlines with angles above 330 degrees).

### Track selection

The full track selection procedure is illustrated in Figure 2. To select tracks for tractometry analyses, connectivity matrices were first created using the Brainnetome atlas, a connectivity-based parcellation with 210 cortical and 36 subcortical labels (Fan et al., 2016). For each subject, a 1mm MNI-152 template (Maintz & Viergever, 1998) was warped onto upsampled DWI space using ANTs SyN algorithm. The Brainnetome atlas is freely available (http://atlas.brainnetome.org/download.html) in MNI-152 space. The resulting transformations from the SyN algorithm were used to convert Brainnetome parcellations onto upsampled DWI space. Using DIPY, a connectivity matrix detailing the number of streamlines connecting each pair of labels was computed (Figure 2A). All streamline counts were normalized by the total amount of streamlines for each subject. The matrices of normalized streamline counts were then binarized using a threshold of at least 1 streamline (Figure 2B).

**Figure 2.**
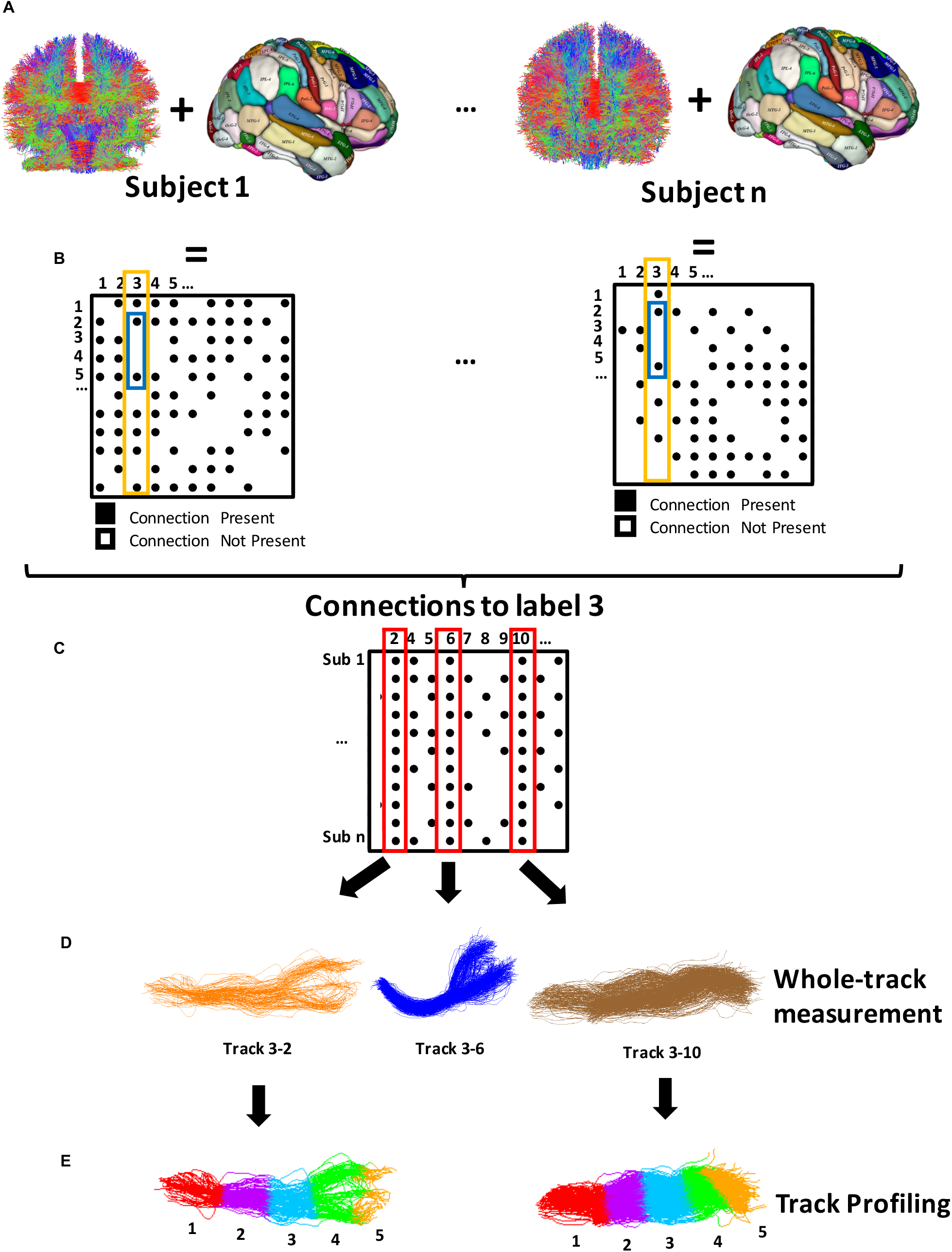
Illustration of the track selection and tractometry methodology. A. Connectomes were built by first segmenting individual whole-brain tractograms using a Brainnetome parcellation (Fan et al. 2016) that was transformed to upsampled DWI space using ANTs. B. Connectomes were binarized such that pairs of regions with at least 1 streamline were assigned a value of 1, and otherwise a value of 0. Labels corresponding to DLPFC regions were selected from binarized connectomes (yellow rectangle), and connections of the DLPFC with other working memory regions were then selected (blue rectangle). C. Matrices containing binarized connectivity values were built for each DLPFC region. These matrices contained all working memory regions selected in the previous step for all subjects. Tracks defined by pairs of regions with an existing connection across all subjects were selected (red rectangle). D. For each selected track, normalized streamline count, normalized track volume, AFD, FW, and FW-corrected FA, AD, MD, RD across were measured across the entire track. E. Tracks in which differences were found underwent track profiling. Tracks were divided in 5 segments, and values of AFD, FW, and FW-corrected FA, AD, MD, RD were measured at each segment.

In the present study, we were interested in tracks connecting the DLPFC with other areas of the brain involved in working-memory. Given that Brainnetome does not contain a DLPFC label, we first defined the DLPFC using 6 labels (Table 1 and Table S1). Next, we defined other regions involved in working memory using 30 labels (Table 1 and Table S1). All labels were selected based on previous literature (Keightley et al., 2014). Tracks that connected one of 6 DLPFC labels with one of 30 working memory labels were isolated (Table 1 and Figure2B). Then, only tracks defined by a working memory label that had an existing connection with a DLPFC label across all 62 subjects were selected for tractometry analyses (Figure 2C). This strict, functionally-informed procedure yielded 11 reliable tracks (see Figure 3).

**Figure 3.**
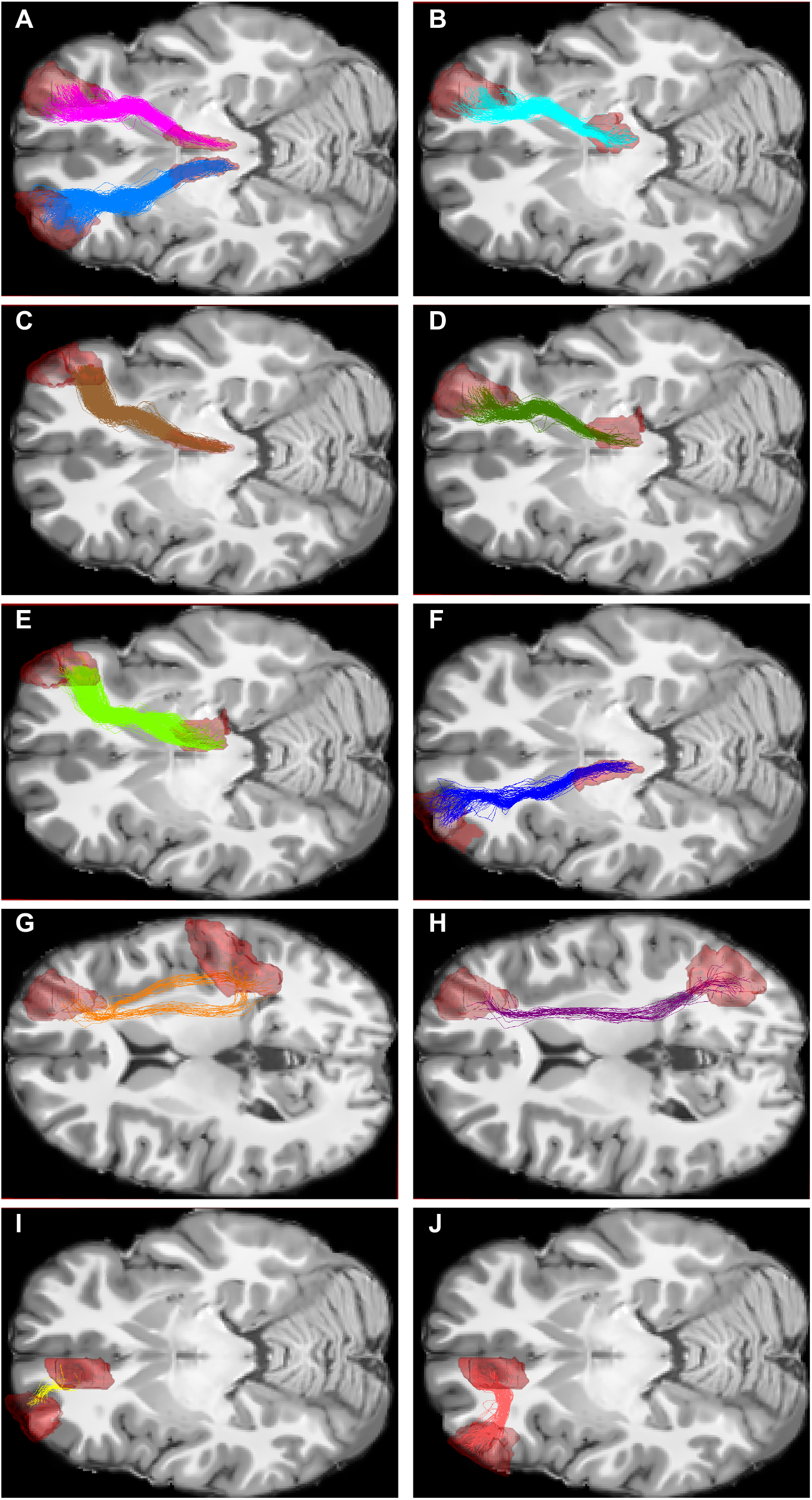
Illustration of the 11 selected working-memory tracks in one subject. End-points (shown in red) are listed in Table 1. A. Magenta: track 16-232; Blue: track 15-231. B: track 16-234. C: track 22-232. D: track 16-246. E: track 22-246. F: track 19-231. G: track 16-140. H: track 16-138. I: track 19-179. J: track 21-179.

**Table 1.** Label numbers and names of endpoints of all studied tracts

| DLPFC Labels |  | Connected Working Memory Areas<sup>a</sup> |  |
| --- | --- | --- | --- |
| Label Number | Label Name<sup>b</sup> | Label Number | Label Name<sup>b</sup> |
| 15 | Left dorsal area 9/46 | 231 | Left medial pre-frontal thalamus |
| 16 | Right dorsal area 9/46 | 138 | Right Inferior Parietal Lobule, rostradorsal area 39 |
|  |  | 140 | Right Inferior Parietal Lobule, rostradorsal area 40 |
|  |  | 232 | Right medial pre-frontal thalamus |
|  |  | 234 | Right pre-motor thalamus |
|  |  | 246 | Right lateral pre-frontal thalamus |
| 19 | Left area 46 | 179 | Left Cingulate Gyrus, pregenual area 32 |
|  |  | 231 | Left medial pre-frontal thalamus |
| 21 | Left ventral area 9/46 | 179 | Left Cingulate Gyrus, pregenual area 32 |
| 22 | Right ventral area 9/46 | 232 | Right medial pre-frontal thalamus |
|  |  | 246 | Right lateral pre-frontal thalamus |

<sup>a</sup>Across 100% of subjects
<sup>b</sup>Anatomical and modified cyto-architectonic descriptions provided with the Brainnetome atlas

### Tractometry

After selecting a reliable set of tracks, we proceeded with a 2-stage track-of-interest analysis. First, the following measurements were taken across the entire track (Figure 2D): normalized streamline count, normalized track volume, FW, Apparent Fiber Density (AFD), and FW-corrected FA, AD, MD, and RD. FW and all FW-corrected tensor-based metrics were extracted from voxel-level measurements, whereas AFD was calculated along streamlines, at the intra-voxel level (Raffelt et al., 2017). Track volumes were normalized by total brain volume. All measurements were weighted by streamline density to minimize the impact of spurious streamlines.

Second, tracks that displayed a significant difference in the first stage were divided into 5 segments (see Figure 2E). The following measurements were taken at each segment in order to create track profiles: FW-corrected FA, AD, MD, RD, as well as AFD and FW. Track profiling was used because it has been shown to be more sensitive than whole-track measurements at detecting subtle changes which correlate well with behaviour (Yeatman, Dougherty, Myall, Wandell, & Feldman, 2012). In follow-up analyses, the Number of Fiber Orientations (NuFO) (Dell’Acqua, Simmons, Williams, & Catani, 2013) were also measured.

### Statistical analyses

Based on skewness and kurtosis analyses of whole-track measurements, streamline count measures were not normally distributed. Hence, group comparisons were performed using Rank-Sum tests on Matlab (Version R2018b), and except where otherwise indicated, group descriptive statistics are reported as medians. Given the known impact of neuropathological processes on diffusion measurements, the following hypotheses were posited. In relation to myelin loss, we hypothesized that FA would be lower in concussed individuals, and RD and MD would be higher (Budde et al., 2011). In relation to axonal loss, we hypothesized that AFD and AD would be lower in concussed individuals (Harsan et al., 2006; Raffelt et al., 2012; S. W. Sun et al., 2006). In relation to edema, we hypothesized that FW would be higher in concussed individuals (Pasternak et al., 2016). Since the objective of the study is to detect these specific patterns of microstructural changes, one-tailed tests according to these hypotheses were used in group comparisons. Given that the direction of change in FA is equivocal (Dodd et al., 2014) in the sub-acute stage, group comparisons of FA, as well as any other comparisons not mentioned above, were performed using two-tailed tests. To assess the impact of microstructure on working memory accuracy, multiple linear regressions were performed using an ordinary least squares estimation method. Regression coefficients are reported in scales that reflect group differences. Regression models were performed using IBM SPSS Statistics for MacIntosh, Version 25.0.

## Results

### Sample after QA

After QA, the selected sample for run 1 consisted of 62 subjects, 16 concussed (10 females, 6 males, mean age=14.75, SD=2.64) and 46 age- and sex-matched healthy controls (17 females, 29 males, mean age=15.09, SD=2.01). Time since injury for the selected concussed participants ranged from 9 to 96 days, with an average of 38.25 days. Despite being balanced for age (CC= 15.80; HC: 15.47; *W*=490, *p*=0.83) and sex (CC= 37.5% males, 62.5% females; HC: 63% males, 37% females; *χ*^2^=3.15, *p*=0.08), groups were not balanced for handedness (CC: 62.5% right handed, 37.5% left handed; HC: 95.7% right handed, 4.3% left handed; χ2=11.61, *p*<0.001). There were no significant differences in age between the included and the excluded participants (included=15.5904, excluded=13.7397, *W*=2306, *p*=0.2040). Information on mechanisms of injury for the included concussed participants is summarized in Table 2.

**Table 2.**
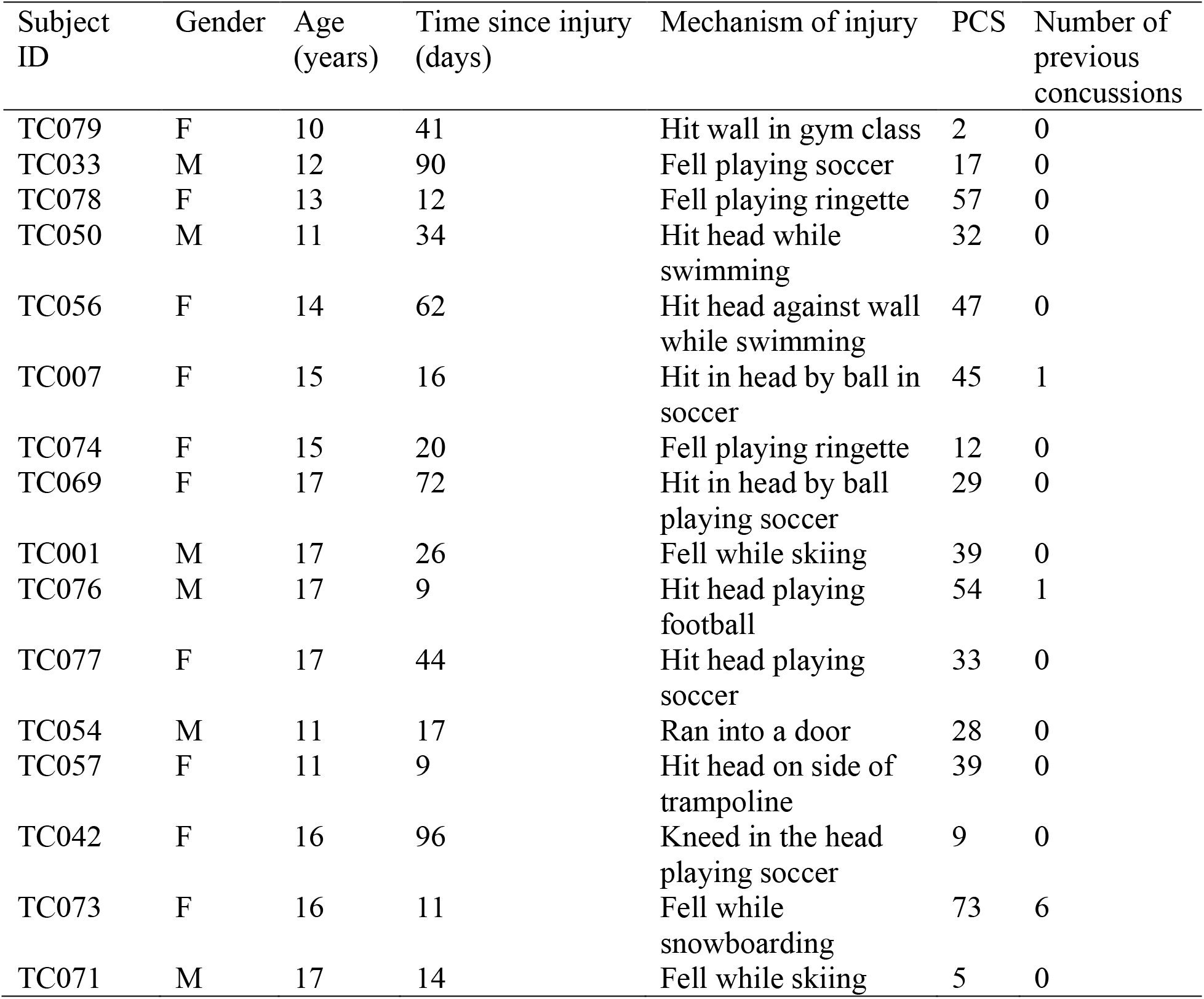
Demographic characteristics of concussed group.

| Subject ID | Gender | Age (years) | Time since injury (days) | Mechanism of injury | PCS | Number of previous concussions |
| --- | --- | --- | --- | --- | --- | --- |
| TC079 | F | 10 | 41 | Hit wall in gym class | 2 | 0 |
| TC033 | M | 12 | 90 | Fell playing soccer | 17 | 0 |
| TC078 | F | 13 | 12 | Fell playing ringette | 57 | 0 |
| TC050 | M | 11 | 34 | Hit head while swimming | 32 | 0 |
| TC056 | F | 14 | 62 | Hit head against wall while swimming | 47 | 0 |
| TC007 | F | 15 | 16 | Hit in head by ball in soccer | 45 | 1 |
| TC074 | F | 15 | 20 | Fell playing ringette | 12 | 0 |
| TC069 | F | 17 | 72 | Hit in head by ball playing soccer | 29 | 0 |
| TC001 | M | 17 | 26 | Fell while skiing | 39 | 0 |
| TC076 | M | 17 | 9 | Hit head playing football | 54 | 1 |
| TC077 | F | 17 | 44 | Hit head playing soccer | 33 | 0 |
| TC054 | M | 11 | 17 | Ran into a door | 28 | 0 |
| TC057 | F | 11 | 9 | Hit head on side of trampoline | 39 | 0 |
| TC042 | F | 16 | 96 | Kneed in the head playing soccer | 9 | 0 |
| TC073 | F | 16 | 11 | Fell while snowboarding | 73 | 6 |
| TC071 | M | 17 | 14 | Fell while skiing | 5 | 0 |

### Tractometry - Whole-track

Our track-selection procedure yielded 11 tracks (Figure 3). The labels that defined each track are outlined in Table 1. From these 11 tracks, two displayed differences in streamline count. Concussed youths had significantly lower streamline count than healthy controls in two thalamo-prefrontal tracks, track 16-246 (Figure 4A, Table 3) and track 22-246 (Figure S1, Table S2). In track 22-246, concussed youths also had lower AD, higher MD, and higher RD (Table S2). Similar differences in AD, MD, and RD were found in another thalamo-prefrontal track, track 22-232, and a cingulo-prefrontal track, track 21-179. Lastly, concussed youths had significantly lower AFD than healthy controls across track 21-179 (Figure5A, Table 4). No significant differences in track volume, FA or FW were apparent across whole-tracks.

**Figure 4.**
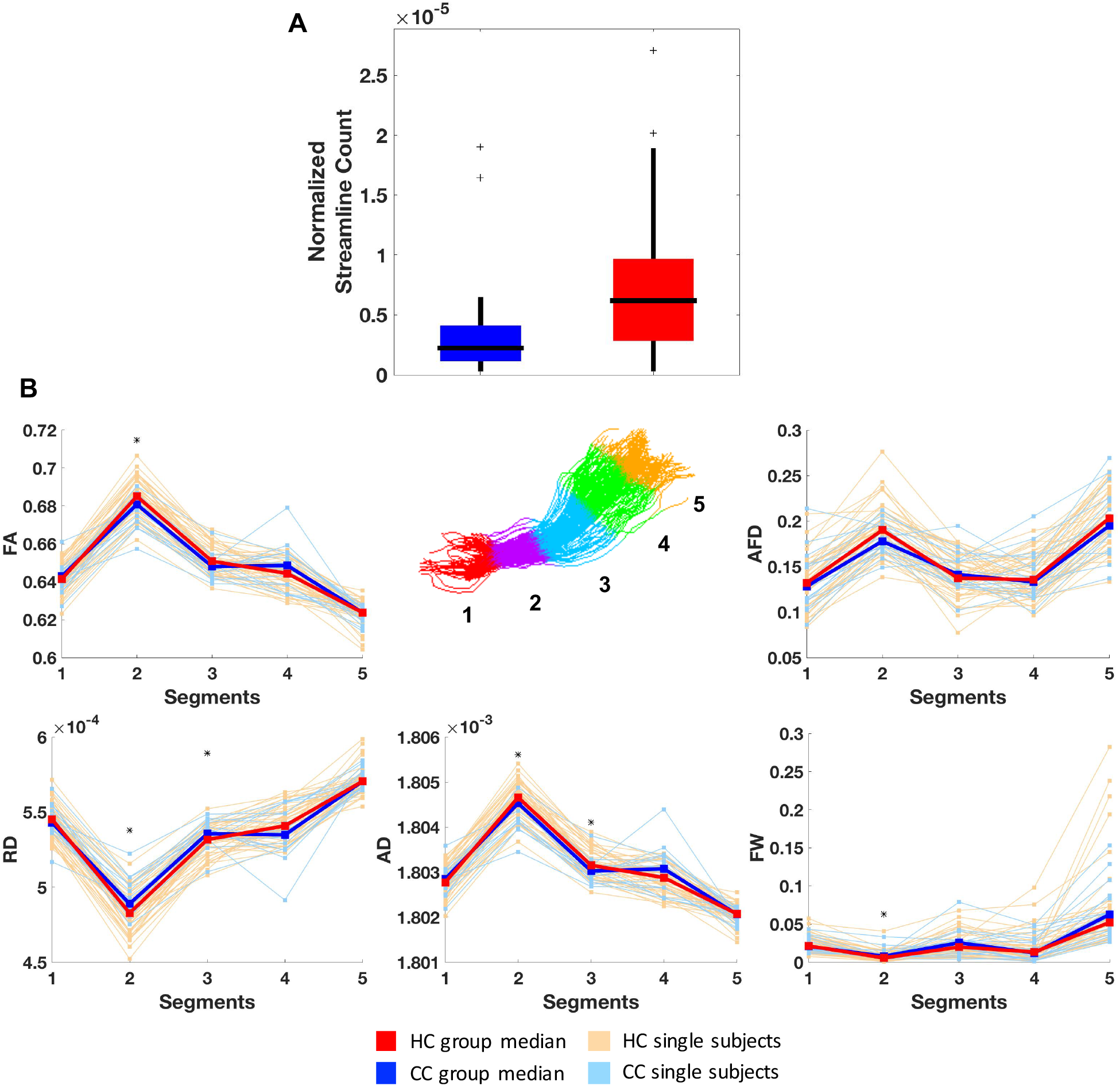
Results of track 16-246. A: Boxplots comparing normalized streamline count between concussed (blue) and healthy control (red) groups. Horizontal black lines illustrate group medians, the edges of the boxes illustrate 75^th^ (upper edge) and 25^th^ (lower edge) percentiles, vertical black lines illustrate ranges, and black ‘+’ signs illustrate outliers. B: Line plots showing track profiles for AFD, FW, and FW-corrected FA, RD, and AD. Light red and light blue lines indicate median values of each segment for each individual healthy or concussed subject respectively. Dark red and dark blue lines indicate instead group medians at each segment. *: *p*<0.05.

**Table 3.** Results for track 16-246

| Segment | Metric | Concussed | Healthy | Mann-Whitney |  |
| --- | --- | --- | --- | --- | --- |
|  |  |  |  | *W* | *p* |
| Whole-track | Streamline Count | $0.223 \cdot 10^{-5}$ | $0.620 \cdot 10^{-5}$ | 346 | 0.0113 |
| 2 | AD | $18.045 \cdot 10^{-4}$ | $18.047 \cdot 10^{-4}$ | 359 | 0.010 |
|  | FA | 0.68 | 0.69 | 357 | 0.018 |
| | MD | $9.275 \cdot 10^{-4}$ | $9.234 \cdot 10^{-4}$ | 651 | 0.009 |
| | RD | $4.890 \cdot 10^{-4}$ | $4.828 \cdot 10^{-4}$ | 651 | 0.009 |
|  | FW | 0.0079 | 0.0057 | 616 | 0.0364 |
| 3 | AD | $18.030 \cdot 10^{-4}$ | $18.032 \cdot 10^{-4}$ | 387 | 0.0305 |
| | MD | $9.581 \cdot 10^{-4}$ | $9.555 \cdot 10^{-4}$ | 614 | 0.039 |
| | RD | $5.357 \cdot 10^{-4}$ | $5.317 \cdot 10^{-4}$ | 614 | 0.039 |

**Table 4.**
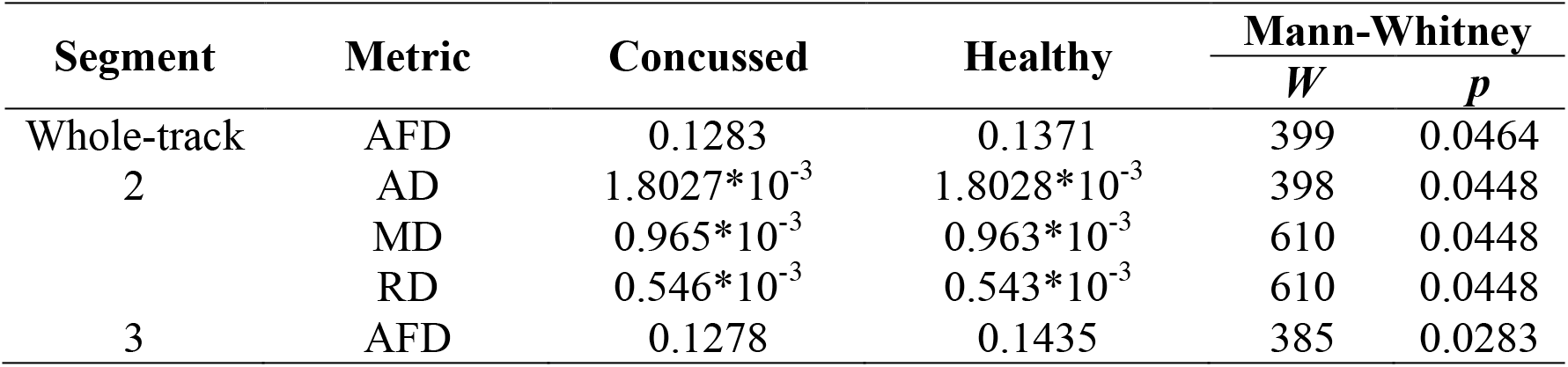
Results for track 21-179

### Tractometry – Track profiling

Track profiles of FA, AD, MD, RD, AFD, and FW were created for tracks 16-246 (Figure 4B, Table 3), 22-246 (Figure S1), 22-232, and 21-179 (Figure 5B, Table 4). Concussed individuals had lower FA than healthy controls in segment 2 of track 16-246. In the same segment, concussed youths had lower AD, higher MD, higher RD, and higher FW than healthy controls. In segment 3 of track 16-246, concussed individuals also displayed lower AD, and higher MD and RD. Concussed individuals also had lower FA than healthy controls in segment 2 of track 22-246, as well as lower AD (Table S2). In the same segment, concussed individuals had higher MD and RD. In segment 3 of this same track, concussed individuals also had lower AD, and higher MD and RD. Concussed individuals also had lower FA than healthy controls in segment 2 of track 22-232, and lower AD. In the same segment, concussed individuals had higher MD and RD. In segment 3 of this same track, concussed individuals also had lower AD, and higher MD and RD (Table S3). Lastly, concussed individuals had lower AFD compared to healthy controls in segment 3 of track 21-179. In segment 2 of track 21-179, concussed individuals had lower AD, higher MD and RD.

**Figure 5.**
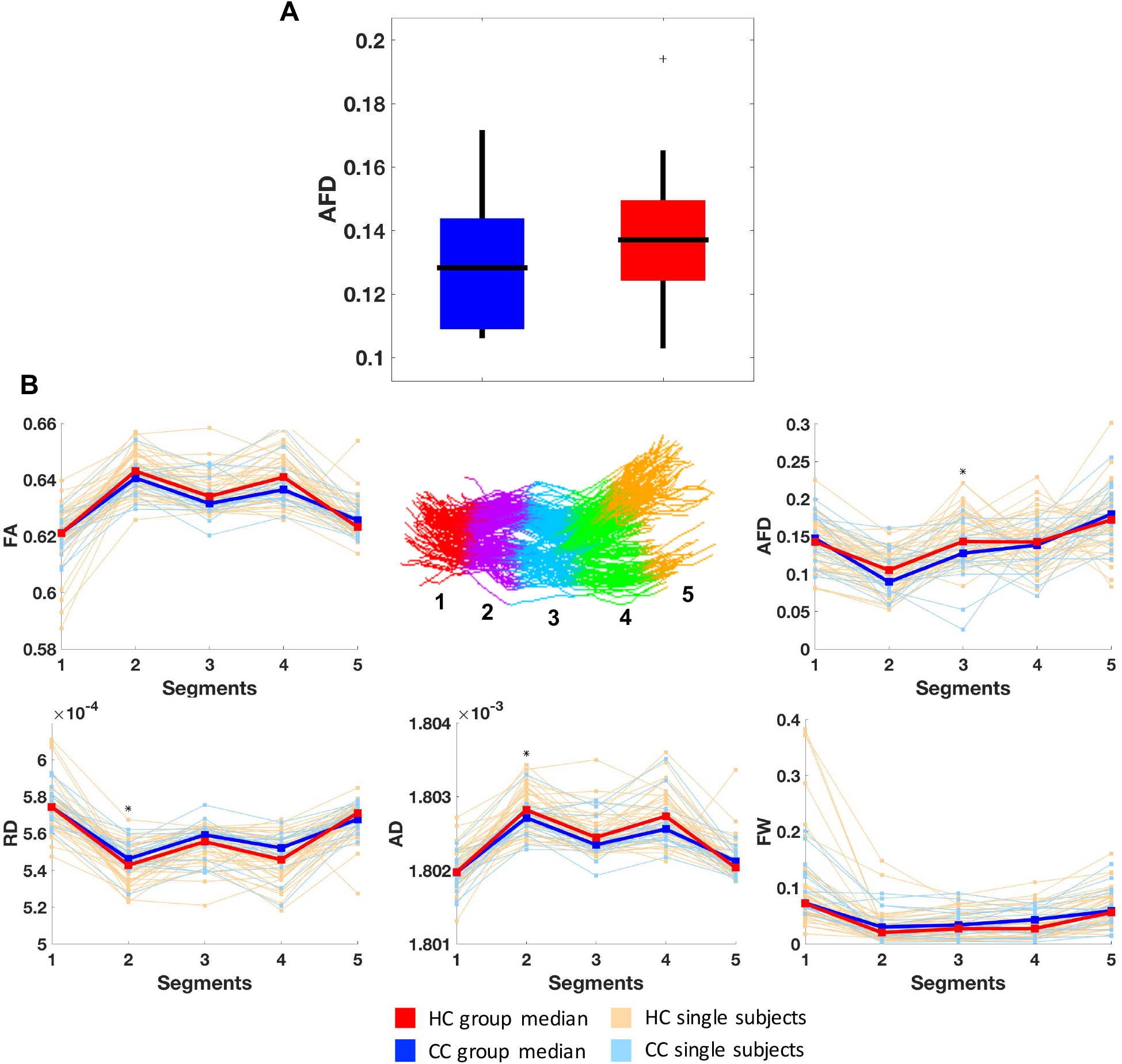
Results of track 21-179. A: Boxplots comparing whole-track AFD between concussed (blue) and healthy control (red) groups. Horizontal black lines illustrate group medians, the edges of the boxes illustrate 75^th^ (upper edge) and 25^th^ (lower edge) percentiles, vertical black lines illustrate ranges, and black ‘+’ signs illustrate outliers. B: Line plots showing track profiles for AFD, FW, and FW-corrected FA, RD, and AD. Light red and light blue lines indicate median values of each segment for each individual healthy or concussed subject respectively. Dark red and dark blue lines indicate instead group medians at each segment. Note: Segment 1 of AD track-profile contained three extreme values that were preventing a proper illustration of the rest of the track profiles. These values were removed for this figure but were kept in analyses. *: *p*<0.05.

### Relation between white-matter and behaviour

Concussed youths had lower verbal (CC: 60, HC: 71; *W*=297, *p*=0.001), and visual working memory accuracy (CC: 58.5, HC: 72; *W*=269, *p*<0.001). To assess whether working memory was associated with white-matter structure, linear regression analyses were computed using white-matter measurements from all significant segments to predict verbal and visual working memory accuracy. Kurtosis and skewness were measured for all track-profile variables that were significantly different between groups. Only one variable, FW in segment 2 of track 16-246, had kurtosis and skewness values suggesting non-normality. This variable was log transformed, which fixed its non-normality. Given that several highly correlated microstructural measurements were entered in the models, backwards step-wise regressions were performed. Once a final set of predictors was obtained, assumptions of linear regression were verified. For track 16-246 (Figure 6A), backwards regression yielded a final model significantly predicting verbal working memory (*F*(1,59)=12.20, *p*=0.001), which accounted for 17% of the variance and contained one significant factor. Participants’ verbal working memory scores increased by 1.097 units for every 1*10^−7^ unit increase of AD in segment 2 of track 16-246. In addition, backwards regression yielded a final model significantly predicting visual working memory (*F*(1,59)=10.43, *p*=0.002), which accounted for 15% of the variance and contained one significant factor. Participants’ visual working memory scores decreased by 0.007 units for every 1*10^−3^-unit increase in the log transformed FW variable in segment 2 of track 16-246 (Figure 5A). Results were similar for tracks 22-246 and 22-232, and are outlined in Supplementary Material.

**Figure 6.**
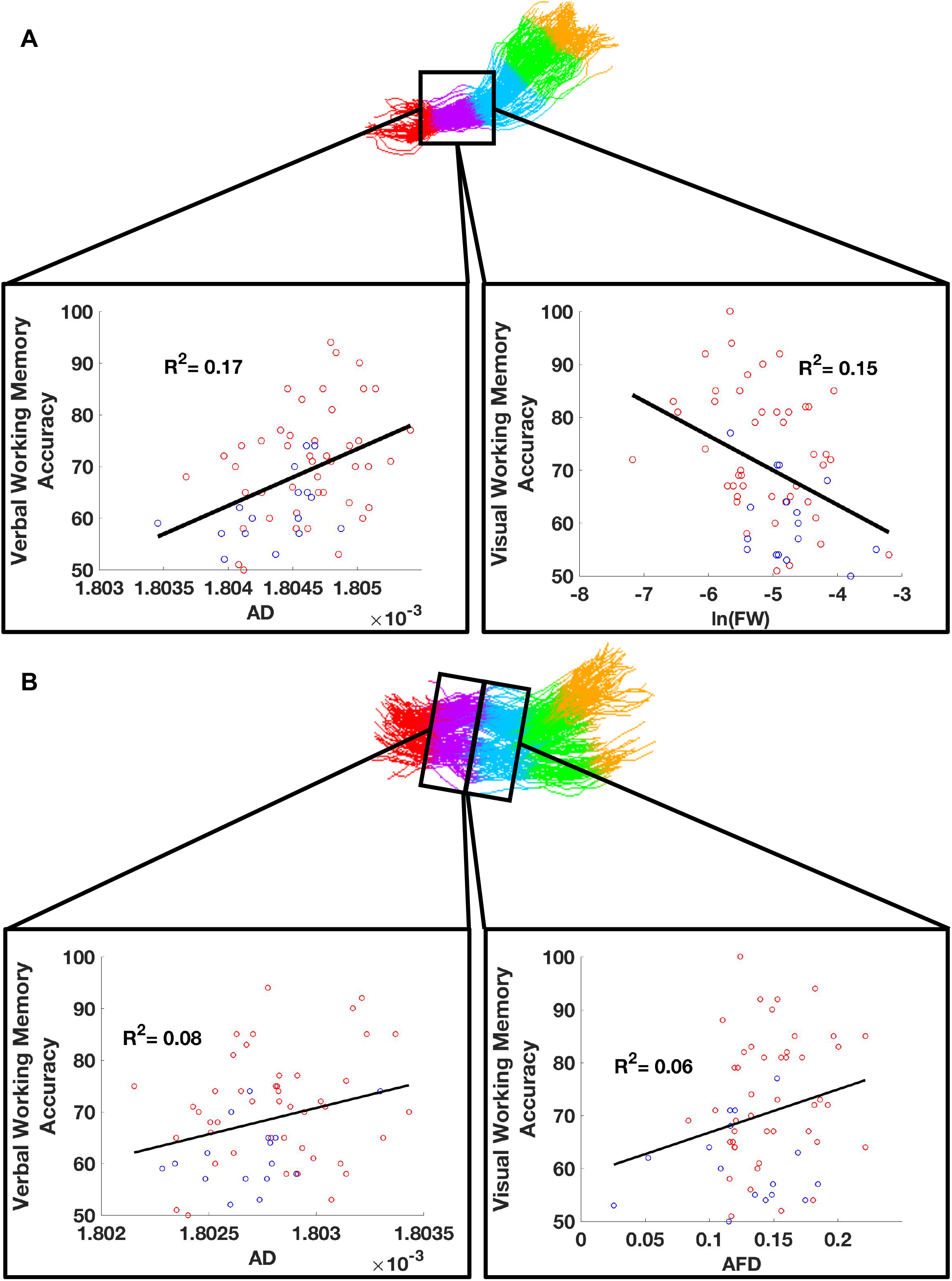
A. Correlations between AD and verbal working memory accuracy (left) and the log-transformed FW and visual working memory accuracy (right) in segment 2 of trakt 16-246. B. Correlations between AD and verbal working memory accuracy (left) in segment 2 and AFD and visual working memory accuracy (right) in segment 3 of track 21-179.

Finally, for track 21-179 (Figure 6B), backwards regression yielded a final model that significantly predicted verbal working memory (F(1,59)=4.979, p=0.029), which accounted for 8% of the variance and contained one significant factor. Participants’ verbal working memory accuracy scores increased 1.020 units for every increase in 1*10^−7^ units of AD in segment 2 of track 21-179. For visual-working memory, backwards regression yielded a final model that did not significantly predict verbal working memory (F(1,59)=3.505, p=0.066.

All assumptions of linear regressions – normality, linearity, homoscedasticity, independence, outliers, and multicollinearity – were verified in the final models and were found to be satisfied.

### Correlation with concussion symptoms

Measures of track microstructure correlated with individual PCS scores in concussed participants (Table 5). In segment 2 of track 16-246, AD was significantly negatively correlated with Remembering Difficulties. In this same segment, log-transformed FW was significantly positively correlated with Vomiting. In segment 2 of track 22-246, FA was significantly negatively correlated with Nausea, and nearly-significantly correlated with Remembering Difficulties. In segment 2 of track 22-232, FA was significantly positively correlated with Nausea and Dizziness. Lastly, in segment 2 of track 21-179, AD was significantly negatively correlated with Noise sensitivity, and nearly-significantly correlated with Remembering Difficulties. Other significant or nearly-significant correlations are listed in Table 5.

**Table 5.**
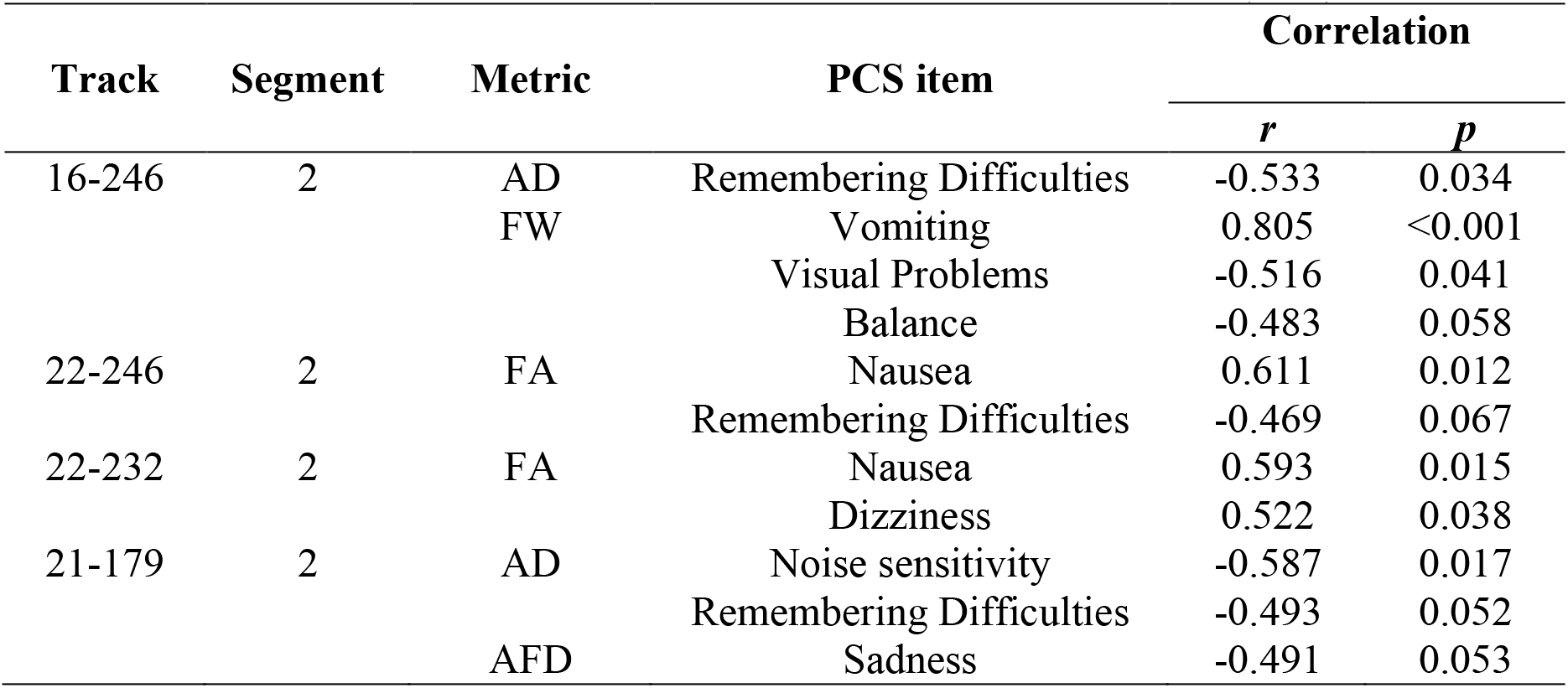
Correlations between track structure and Post-Concussion Scale (PCS) items

### Number of Fiber Orientations

To assess whether changes in FA, AD, MD, and RD of tracks 16-246, 22-246, and 22-232 were due to changes in the number of crossing fiber populations, NuFO values were computed along these tracks. No significant differences were found between groups.

### Replication on run 2

To assess the reliability of the present results, analyses that yielded significant group differences with run 1 data were repeated with run 2 data. For the subset of individuals whose second scans passed initial QA (n=40, 10 concussed, 30 controls), we first selected tracks using the same criteria as in run 1. Ten tracks were selected from run 2 scans, only 2 of which were not within the run 1 tracks. From the four tracks that showed tractometry differences in run 1, three (16-246, 22-232, and 21-179) were among the 10 selected from run 2 scans. Only one healthy control did not have a streamline for track 22-246. Nonetheless, only results obtained in tracks 16-246, 22-232, and 21-179 were considered for this replication phase.

In run 2 scans (Table S4), concussed youths had a significantly lower normalized streamline count than healthy controls in track 16-246, a lower AD, higher MD and RD across track 21-179. In addition, in run 2 scans, concussed youths also had lower AD, higher MD and RD in track 16-246 segment 2, track 22-232 segments 2 and 3, and track 21-179 segment 2. Lastly, concussed youths had lower AFD in segment 3 of track 21-179.

### Voxel Overlap Scores

On the subset of individuals whose second scans passed initial QA, voxel-overlap scores were computed to assess the reliability of our tractography procedure. FA maps from run 2 scans were converted into run 1 DWI space using ANTs. Overlap scores were computed by assessing the amount of voxels traversed by the same track in both scans, and scores were weighted by track density to minimize the effect of spurious tracks. Across 40 participants, track 16-246 had an average voxel-overlap score of 0.81 (SD=0.09), track 22-232 an average of 0.85 (SD=0.08), and track 21-179 an average of 0.66 (SD=0.16).

### Total number of streamlines

The total number of streamlines was variable (SD=7.58*10^5^). Although streamline counts for each track were normalized by total streamline count for each subject, we wanted to ensure that concussed individuals did not have a feature that made tracking globally harder. We thus compared total streamline counts between groups. No significant differences were found (CC: 1.98*10^6^, HC: 2.06*10^6^; *W*=446, *p*=0.3550). If fiber tracking is more difficult in concussed individuals, we would also expect reconstructions to be less reliable. Hence, we compared voxel-overlap scores for the three tracks of interest between groups. No significant differences were seen for track 16-246 (CC: 0.793, HC: 0.837; *W*=182, *p*=0.488), 22-232 (CC: 0.86, HC: 0.87, *W*=180, *p*=0.22), or 21-179 (CC: 0.603, HC: 0.690; *W*=116, *p*=0.233).

## Discussion

In the present study, we leveraged for the first time recent advances in DWI local modelling (CSD, free-water modelling), tractography (PFT with anatomical priors), and tractometry (AFD, FW, FW-corrected tensor measures, and track profiling), to better understand the neuropathological basis of working-memory deficits after concussions. After a rigorous, functionally-informed, track-selection procedure, we identified 11 reliable working-memory tracks. For four of these tracks, we found distinct patterns of microstructural changes, which were correlated to different degrees with working memory performance and with patient-reported memory impairments. Structural differences and tractography reconstructions were reproduced using test-retest analyses.

### Interpreting microstructural differences at whole-track and segment-levels

At the whole-track level, concussed youths displayed lower streamline count for two thalamo-prefrontal tracks, lower AD and higher RD and MD for three thalamo-prefrontal tracks and one cingulo-prefrontal track, and lower AFD in the cingulo-prefrontal track. At the segment-level, concussed youth also demonstrated lower FA and AD, higher RD and MD in the same segment of the three thalamo-prefrontal tracks. Animal studies suggest that decreases in FA accompanied by increases in RD and MD are suggestive of alterations in myelin structure (Budde et al., 2011). Further, an animal model of axonal damage and myelin loss in the corpus callosum demonstrated that decreases in AD are specific to axonal loss (S. W. Sun et al., 2006). However, DTI analyses in this study were performed in the middle section of the corpus callosum, where fibers are homogenously oriented. Given that crossing fibers reduce the anisotropy of the overall voxel, we speculate that the link between AD and axonal density should theoretically decrease when the amount of crossing fibers increases. In contrast, AFD can provide information about axonal density along fascicles, which will be less affected by crossing fibers (Raffelt et al., 2012). In the present study, it is unclear whether axonal loss is present in the thalamo-prefrontal tracks, given that AFD and NuFO were not significantly changed (Jones et al., 2013). A study on mouse models of mTBI examined progression of white matter pathology and found a pattern of acute myelin loss at 3 days followed by a pattern of remyelination at 1 week and excessive myelination by 6 weeks (Mierzwa, Marion, Sullivan, McDaniel, & Armstrong, 2015). In a recent study, Spader et al. (2018) used multicomponent driven equilibrium single pulse observation of T1 and T2, a quantitative myelin imaging technique, to study 27 collegiate athletes. They found that athletes had higher myelin water fraction (MWF), a measure of myelin content, when compared to healthy controls at 72 hours after injury. In addition, concussed athletes had higher MWF after 3 months compared to themselves at 72 hours, supporting the interpretation that an active remyelination process takes place in the brain after concussions. The patterns of change observed in thalamo-prefrontal tracks in concussed youths in the present study are therefore consistent with an altered, possibly excessive and disordered, myelin structure.

In the cingulo-prefrontal track, the patterns of change, which included higher RD and MD were also suggestive of myelin structure alterations. However, AD and AFD were lower in concussed youths in this track, which is more suggestive of axonal loss. Patterns of tensor-based changes suggestive of axonal loss have been previously observed in studies of concussed individuals (Goswami et al., 2016; Mac Donald et al., 2011). The present study confirms such findings by demonstrating differences in AFD (Raffelt et al., 2012). The presence of potential axonal loss in concussed individuals is important, as these injuries are often believed to be transient (Sharp & Jenkins, 2015).

Only one segment of track 16-246 was found to have increased FW in concussed individuals, and this finding was not replicated in run 2 scans. Although edema is an important and well-documented concussive neuropathology (Bigler & Maxwell, 2012; Hayes et al., 2016; Werner & Engelhard, 2007), the present results suggest that this pathology was not widespread in this particular concussed group. Animal models of TBI suggest that edema is resorbed after 1 week (Bareyre, Wahl, McIntosh, & Stutzmann, 1997), so perhaps it had been nearly completely resolved in our participants. The group difference in FW reported in the present study was not replicated in test-retest analyses, suggesting that it may possibly be an artefact. This interpretation is also suggested by the paradoxical negative correlations between FW and PCS scores for Balance and Visual Problems. On the other hand, the model using FW to predict visual working memory accounted for 15% of the variance, and FW was highly, positively correlated to the PCS score for Vomiting (*r*=0.805, *p*<0.001). The interpretation for these results is therefore unclear.

Lastly, based on regression analyses, only measures of the thalamo-prefrontal tracks were significantly related to verbal and visual working memory accuracy. White matter structure in the cingulo-prefrontal track was either weakly related to verbal working memory, or unrelated to visual working memory. These results suggest that neuropathology, in this case suggestive of myelin structure alterations, in thalamo-prefrontal tracks appears to play a key role in working memory deficits in concussion. Consistent with the present results, a recent study performed in mice found that thalamus-to-PFC circuits were important for supporting working memory maintenance (Bolkan et al., 2017). Previous studies have repeatedly found that damage to the thalamus plays an important role in the development of cognitive dysfunctions after mTBIs (Abdel-Dayem et al., 1998; Anderson, Wood, Bigler, & Blatter, 1996; Ge et al., 2009; Grossman et al., 2012; Grossman & Inglese, 2016; Henninger et al., 2007; Little et al., 2010; Wood & Bigler, 1995). Further, in fMRI studies, the thalamus has been repeatedly reported among areas demonstrating abnormal activity in concussed individuals (Chen et al., 2007; Chen et al., 2004; Jantzen et al., 2004; Keightley et al., 2014; McAllister et al., 1999; McAllister et al., 2001; Pardini et al., 2010; Westfall et al., 2015). Out of 21 PCS items, symptom scores for Remembering Difficulties were among the few that were moderately (*r*=0.4-0.6) correlated with microstructural measurements of the studied tracks. Significance was only reached for track 16-246, potentially due to small sample-size. AD in track 21-179 was also negatively correlated with self-reported Remembering Difficulties (*r*=-0.493, *p*=0.052), albeit non-significantly. One potential explanation for this result is that PCS ratings for Remembering Difficulties may encompass more general memory function than the presently-used working-memory task.

It is unclear whether these neuropathological findings are generalizable. It is possible that due to demographics or the mechanism of injury in our group (sports injuries), or another unaccounted factor, thalamo-prefrontal tracks may be most affected. However, simulation studies suggest that, given its position near the base of the brain where space is more restricted, the thalamus is among the regions that experiences the highest shear strain from forces transmitted during various forms of head injury mechanisms (Bayly et al., 2005; Sabet, Christoforou, Zatlin, Genin, & Bayly, 2008; Viano et al., 2005; L. Zhang, Yang, & King, 2004). It is therefore possible that these thalamo-frontal tracks tend to be affected across mTBI subjects, irrespective of the mechanism of injury. Factors specific to the present dataset may also possibly explain why the studied thalamo-prefrontal tracks predominantly display a demyelinating profile. Although an imperfect remyelination mechanism may explain why some patients recover from symptoms while others do not, a recent study convincingly showed that in the transneuronal spread of neurodegenerative diseases, axonal loss may always be preceded by myelin alterations, suggesting that neuroinflammation may be playing a role (You et al., 2019). Given the known neuroinflammatory response in concussions, the different neuropathologies observed may instead be parts of a continuum rather than parallel pathologies.

In the present study, we found patterns of change in segments of tracks that were not found at the whole-track level. Across the three thalamo-prefrontal tracks, the same segment, (segment 2 starting from the thalamus) tended to show differences between groups. Two of these tracks shared the same origin in the thalamus and only diverged near the cortex. The other track had a different thalamic origin but shared the same prefrontal termination as one of the other tracks. Given the proximity of all the tracks, it is possible that the pattern of change arose from damage to a single white-matter tract, the thalamic-prefrontal-peduncle (C. Sun et al., 2018). Localized changes in white-matter structure can be influenced by the shape of the tract at different segments or the proximity of other structures (Yeatman et al., 2012). Although fiber crossings at different parts of these tracks are an unlikely explanation, it is nonetheless unclear what exact factors contributed to the highly localized changes observed in the present study.

### Technical considerations

This study is the first to apply CSD and other recent advances in diffusion-MRI to study pediatric concussions. A recent study applied another novelty of diffusion-MRI, the Neurite Orientation Dispersion and Density Imaging (NODDI) model, to study concussions in a sample of individuals aged 18 to 55 (Palacios et al., 2018). They found decreases in FA and MD early after injury, as well as elevated free water fraction. They also found longer-term declines in neurite density. The authors attributed the early changes to vasogenic edema and the later changes to neuronal loss. These results also demonstrate the presence of distinct neuropathologies in concussed individuals, although notable differences exist between the two studies. First, Palacios et al. (2018) studied adults, whereas we studied children. This difference is important given that mTBI incidences peak in children and adolescents (Faul et al., 2010), concussed youth have worse outcomes than adults (Hessen, Nestvold, & Anderson, 2007; McKinlay, Corrigan, Horwood, & Fergusson, 2014), and far fewer biomarker studies exist in children than adults (Mayer et al., 2018). Second, Palacios et al. (2018) used NODDI, whereas we used CSD, novel tractography methods, FW-corrections, and track profiling to help refine track segmentation and measurements of white-matter microstructure. This difference is important because, despite its advantages over a classical tensor model, NODDI still assumes a single-fiber population, whereas the methods used presently are more robust to this assumption, especially CSD, which can account for crossing fibers better than NODDI. The addition of more specific measurements of white matter microstructure (FW, NuFO, AFD) helped complement the information provided by tensor measures. Nonetheless, these tensor-based measurements remain voxel-level measurements and thus not track-specific. Instead, CSD gives rise to fODFs and fiber elements (or “fixels”) (Raffelt et al., 2017) which can provide intra-voxel, orientation-dependent fiber density information along fiber fascicles. AFD computed along “fixels” thus provided track-specific microstructural measurements that are more neuropathologically-informative. Another important methodological contrast is that NODDI requires specialized acquisition protocols that may not be available outside large urban hospital centers, whereas the tools used in this study can theoretically be applied to any existing dataset. Although HARDI data is optimal for CSD, the anatomical accuracy of tractography reconstructions can be drastically improved with CSD even when using suboptimal acquisition protocols (Farquharson et al., 2013), which may be the only available in more remote settings. It is reasonable to contend that a criterion for a clinically useful biomarker is one that can be made widely available, and in that regard, the present approach may be more clinically promising than the one used in the Palacios et al. (2018) study.

### Limitations and Strengths

First, groups were not balanced for handedness. In a study of 42 right- and 40 lefthanders aged 18 to 31, Powell et al. (2012) used voxel-based statistics and found that selfreported right-handers had higher FA in the left hemisphere’s limbic cortex, PFC, medial frontal lobe, and inferior frontal gyrus, and the right hemisphere’s orbitofrontal cortex, middle and inferior frontal gyri. It is possible that our concussed group, which contained significantly more left-handers, had lower FA than healthy controls because of handedness rather than injury. Powell et al. (2012) correctly argue that their results depend on an assumption of similar numbers of crossing fibers between left- and right-handers. No conclusive evidence exists to support this assumption, and the present study, by using new methods that can provide orientation-specific information that is robust to underlying fiber configuration, can overcome the potentially confounding factors that may be present in the Powell et al. (2012) study. Second, the sample size was relatively small for the number of tests performed, raising concerns about insufficiency of power. Several efforts were taken to limit the number of tests, which, when using track-profiling, could have been exponentially greater, but as a consequence, other important differences may have been missed. Replication with larger datasets is necessary. Although the neuropathological interpretations advanced in this study are supported by previous and recent literature, these techniques do not offer the same level of specificity as other forms of microstructural imaging. Replication with other modalities, especially with myelin-based contrast imaging, and validation with histological studies are therefore still needed. Lastly, the cross-sectional design of this study prevents inferences on clinical outcomes. Future longitudinal studies of concussed children are needed to assess whether patterns of change suggesting different types of neuropathologies are associated with different clinical outcomes.

In addition to using novel methodology, strengths of the present study include the use of a sample of age- and sex-matched controls, a high-angular resolution dataset to ensure rotationally-invariant tensor (Jones, 2004) and fODF reconstructions (Jones et al., 2013), and a strict QA procedure. Further, the use of strict track-selection criteria ensured that tractometry analyses were performed only on a reliable set of tracks. The tracks with the majority of effects, 16-246, 22-232, and 21-179, were obtained across all 102 selected scans, making these tracks highly replicable. Voxel-overlap scores between run 1 and run 2 tracks also reflect this high reliability. Although the average voxel-overlap score in track 21-179 was lower than track 16-246, its standard deviation was larger, suggesting that this track may be harder to reconstruct, perhaps due to its particular anatomical location and proximity to major white-matter bundles (Figure S2). Results on this track were replicated on run 2 data, despite the higher difficulty associated with its reconstruction. Lastly, the use of PCS scores provided a way to validate the clinical relevance of track measurements.

### Conclusion and clinical considerations

This study is the first to apply major recent advances in diffusion-MRI to better understand the neuropathology involved in working memory deficits in sub-acutely concussed youths. The present results suggest that myelin structure alterations in thalamo-prefrontal tracks play a key role in working-memory impairments after concussions. More broadly, this study is the first to observe patterns suggesting the presence of concurrent pathologies in the brains of concussed children. Although no universal consensus exists for the definition of a biomarker, prior work suggests that a clinically useful biomarker needs to reliably and objectively detect the presence or absence of disease, track the rate of disease progression, and provide information on the mechanism of disease and clinical outcome in order to inform the most appropriate course of therapy (Holland, 2016; Salter & Holland, 2014). This last criterion is often overlooked in concussions. Clinical outcomes of concussed patients vary considerably, and symptomatic pharmacological therapy is prescribed using a “wait and see” approach, whereby treatment is only given when symptoms become chronic (above 3 months) (Broglio, Collins, Williams, Mucha, & Kontos, 2015). In contrast, emerging research suggests that early pharmacotherapy may be more effective, especially when tailored to the specific clinical profile of the patient (Broglio et al., 2015). Hence, neuropathologically-informed, clinically-useful biomarkers are still needed, and by providing a better understanding of the neuropathology involved in sub-acute pediatric concussions, the present study constitutes an important first step.

## Acknowledgements

We want to thank participants and their parents for lending their time for this study. We also want to thank Dr. Jen-Kai Chen and Dr. Rajeet Singh Salujah for their contribution in the acquisition of this data.

