## Supplementary material for "Structural abnormalities in thalamo-prefrontal tracks revealed by high angular resolution diffusion imaging predict working memory scores in concussed children": Figure S1

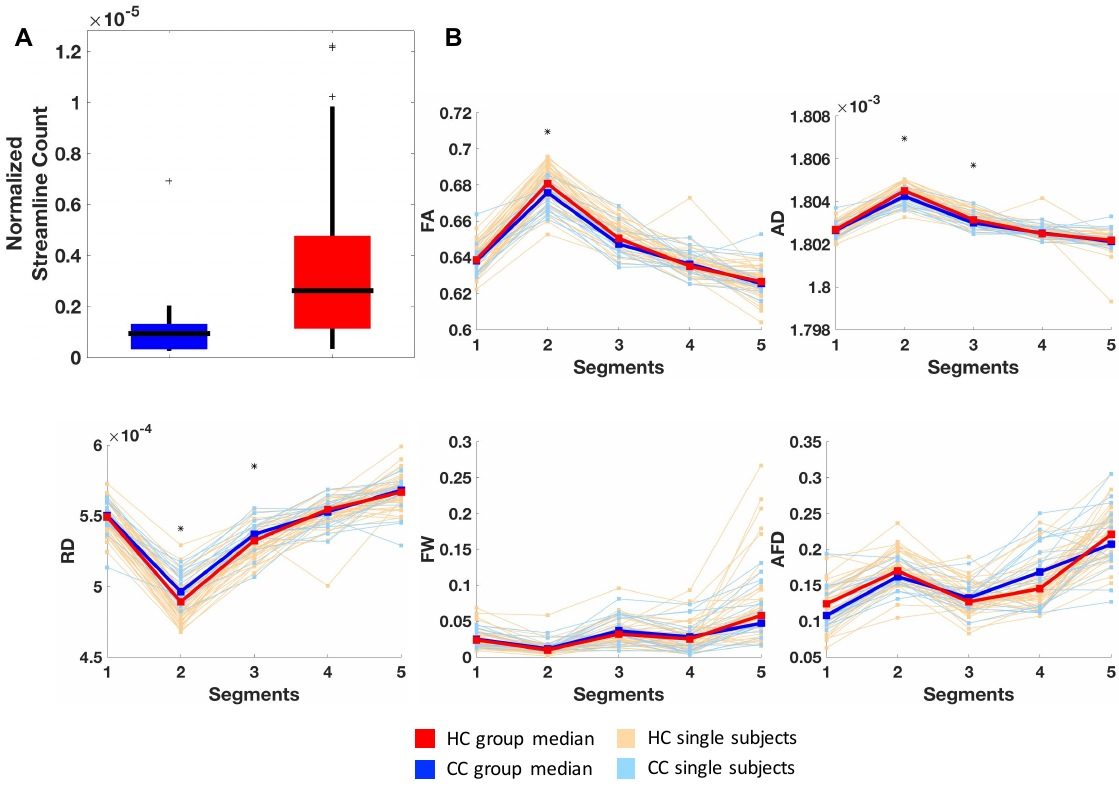

Figure S1. Results of track 22-246. A: Boxplots comparing normalized streamline count between concussed (blue) and healthy control (red) groups. Horizontal black lines illustrate group medians, the edges of the boxes illustrate 75^th^ (upper edge) and 25^th^ (lower edge) percentiles, vertical black lines illustrate ranges, and black ‘+’ signs illustrate outliers. B: Line plots showing track profiles for AFD, FW, and FW-corrected FA, RD, and AD. Light red and light blue lines indicate median values of each segment for each individual healthy or concussed subject respectively. Dark red and dark blue lines indicate instead group medians at each segment. *: *p*<0.05.
