## Supplementary material for "Structural abnormalities in thalamo-prefrontal tracks revealed by high angular resolution diffusion imaging predict working memory scores in concussed children": Figure S2

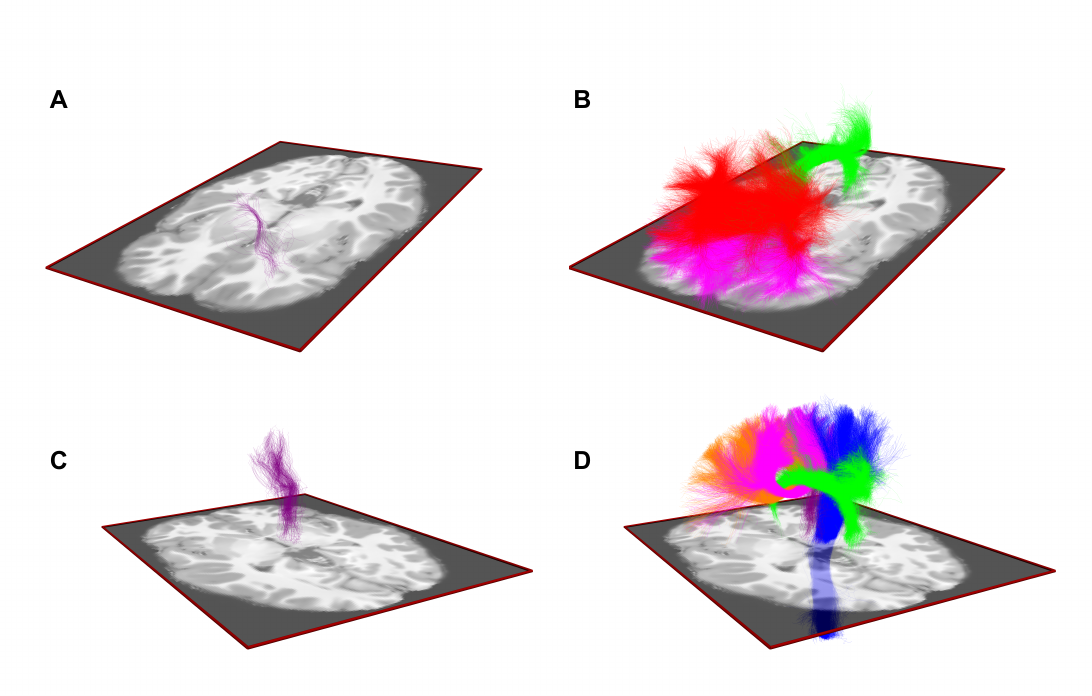

Figure S2. A. Illustration of track 21-179 for one subject, without other tracks visible. B. Illustration of track 21-179 for the same subject, with the cingulum (green) and two segments of the corpus callosum (magenta and red). Track 21-179 is completely occluded by these large tracks. C. Illustration of track 16-246 for one subject, without other tracks visible. D. Illustration of track 16-246 for the same subject, with cingulum (green), corticospinal track (blue), and two other segments of the corpus callosum (magenta and orange). Track 16-246 is still visible.
