## Supplementary Material for "Structural abnormalities in thalamo-prefrontal tracks revealed by high angular resolution diffusion imaging predict working memory scores in concussed children"

### *Quality Assessment*

All diffusion images underwent a rigorous quality assessment procedure. First, all raw diffusion images were visually inspected and rated on a scale from 0 to 7 based on the presence (score of 0) or absence (score of 1) of the following 7 artifacts: motion, noise, distortions (e.g.: EPI distortion), truncations, slicing and spike drops, magnetic field inhomogeneity, and ghosting. Total scores were summed and used to categorize images as either Poor (score of 0-3), Good (4-5), or Excellent (6-7). Images rated as Poor were flagged for removal. In parallel, an automated quality assessment algorithm, described in Roalf et al. (2016), was also implemented. Following their recommendations, total signal-to-noise ratio was computed and images with a tSNR below 6.47 were flagged for removal. This process allowed us to obtain two individual ratings, one from a visual rater, and one from the automatic algorithm. These two ratings had an agreement of 82% across the 71 run 1 scans. For subjects whose run 1 data was excluded on the basis of this QA procedure, run 2 scans were also excluded. The run 2 scans for the remaining subjects also underwent the same QA procedure. Agreement for run 2 scans was of 61% between both raters. Images that were in agreement were either definitely removed or definitely kept. Images in which the two ratings disagreed were flagged for follow-up after processing. A second visual inspection of the processed images which were initially flagged for follow-up, performed after preprocessing, determined whether or not these images would be used for the next phases.

### *Results for regression analyses*

For track 22-246, backwards regression yielded a final model significantly predicting verbal working memory ($F(1,59)=15.556$, $p<0.001$), which accounted for 21% of the variance and contained one significant factor. Participants' verbal working memory accuracy scores increased 0.477 units for every $1*10^{-3}$-unit increase of FA in segment 2 of track 22-246. For this same track, backwards regression yielded a final model significantly predicting visual working memory ($F(1,59)=7.176$, $p=0.010$), which accounted for 11% of the variance and contained one significant factor. Participants' visual working memory accuracy scores increased 0.406 units for every $1*10^{-3}$-unit increase of FA in segment 2 of track 22-246.

For track 22-232, backwards regression yielded a final model significantly predicting verbal working memory ($F(1,59)=13.568$, $p=0.001$), which accounted for 19% of the variance and contained one significant factor. Participants' verbal working memory accuracy scores increased 0.455 units for every increase in $1*10^{-3}$ units of FA in segment 2 of track 22-232. For this same track, backwards regression yielded a final model significantly predicting visual working memory ($F(1,59)=7.491$, $p=0.008$), which accounted for 11% of the variance and contained one significant factor. Participants' visual working memory accuracy scores increased 0.417 units for every $1*10^{-3}$-unit increase of FA in segment 2 of track 22-232.

Table S1. List of label numbers and names of endpoints that were not part of included tracks

| DLPFC Labels | | Non-Connected Working Memory Areas[a] | |
|---|---|---|---|
| Label Number | Label Name[b] | Label Number | Label Name[b] |
| 20 | Right area 46 | 127 | Left Superior Parietal Lobule, caudal area 7 |
| | | 128 | Right Superior Parietal Lobule, caudal area 7 |
| | | 129 | Left Superior Parietal Lobule, lateral area 5 |
| | | 130 | Right Superior Parietal Lobule, lateral area 5 |
| | | 133 | Left Superior Parietal Lobule, intraparietal area 7 |
| | | 134 | Right Superior Parietal Lobule, intraparietal area 7 |
| | | 137 | Left Inferior Parietal Lobule, rostrodorsal area 39 |
| | | 139 | Left Inferior Parietal Lobule, rostrodorsal area 40 |
| | | 180 | Right Cingulate Gyrus, pregenual area 32 |
| | | 183 | Left Cingulate Gyrus, caudodorsal area 24 |
| | | 184 | Right Cingulate Gyrus, caudodorsal area 24 |
| | | 233 | Left pre-motor thalamus |
| | | 235 | Left sensory thalamus |
| | | 236 | Right sensory thalamus |
| | | 237 | Left rostral temporal thalamus |
| | | 238 | Right rostral temporal thalamus |
| | | 239 | Left posterior parietal thalamus |
| | | 240 | Right posterior parietal thalamus |
| | | 241 | Left occipital thalamus |
| | | 242 | Right occipital thalamus |
| | | 243 | Left caudal temporal thalamus |
| | | 244 | Right caudal temporal thalamus |
| | | 245 | Left lateral pre-frontal thalamus |

[a]Across 100% of subjects

[b]Anatomical and modified cyto-architectonic descriptions provided with the Brainnetome atlas

Table S2. Results for track 22-246

| Segment | Metric | Concussed | Healthy | Mann-Whitney | |
|---|---|---|---|---|---|
| | | | | ***W*** | ***p*** |
| Whole-track | Streamline Count | $0.0933*10^{-5}$ | $0.261*10^{-5}$ | 270 | <0.001 |
| | AD | $18.03*10^{-4}$ | $18.03*10^{-4}$ | 397 | 0.043 |
| | MD | $9.56*10^{-4}$ | $9.53*10^{-4}$ | 611.5 | 0.043 |
| | RD | $5.32*10^{-4}$ | $5.28*10^{-4}$ | 611 | 0.043 |
| 2 | AD | $1.8042*10^{-3}$ | $1.8045*10^{-3}$ | 349 | 0.0065 |
| | FA | 0.6757 | 0.6808 | 359 | 0.0201 |
| | MD | $0.9322*10^{-3}$ | $0.9275*10^{-3}$ | 649 | 0.0100 |
| | RD | $0.4962*10^{-3}$ | $0.4890*10^{-3}$ | 649 | 0.0100 |
| 3 | AD | $1.8030*10^{-3}$ | $1.8031*10^{-3}$ | 389 | 0.0327 |
| | MD | $0.9588*10^{-3}$ | $0.9599*10^{-3}$ | 617 | 0.0352 |
| | RD | $0.5367*10^{-3}$ | $0.5322*10^{-3}$ | 617 | 0.0352 |

Table S3. Results for track 22-232

| **Segment** | **Metric** | **Concussed** | **Healthy** | **Mann-Whitney** | |
|---|---|---|---|---|---|
| | | | | ***W*** | ***p*** |
| 2 | AD | $1.8040*10^{-3}$ | $1.8043*10^{-3}$ | 353 | 0.0077 |
| | FA | 0.6706 | 0.6761 | 354 | 0.0162 |
| | MD | $0.9370*10^{-3}$ | $0.9319*10^{-3}$ | 654 | 0.0081 |
| | RD | $0.5035*10^{-3}$ | $0.4957*10^{-3}$ | 654 | 0.0081 |
| 3 | AD | $1.8030*10^{-3}$ | $1.8031*10^{-3}$ | 400 | 0.0480 |
| | MD | $0.9594*10^{-3}$ | $0.9568*10^{-3}$ | 607 | 0.0496 |
| | RD | $0.5376*10^{-3}$ | $0.5336*10^{-3}$ | 607 | 0.0496 |

Table S4. Results for run 2

| **Track** | **Segment** | **Metric** | **Concussed** | **Healthy** | **Mann-Whitney** | |
|---|---|---|---|---|---|---|
| | | | | | ***W*** | ***p*** |
| 16-246 | Whole-track | Streamline Count | $2.74*10^{-6}$ | $5.71*10^{-6}$ | 128 | 0.008 |
| | 2 | AD | $1.8042*10^{-3}$ | $1.8044*10^{-3}$ | 151 | 0.0474 |
| | | MD | $0.9332*10^{-3}$ | $0.9293*10^{-3}$ | 259 | 0.0474 |
| | | RD | $0.4976*10^{-3}$ | $0.4918*10^{-3}$ | 259 | 0.0474 |
| 21-179 | Whole-track | AD | $1.8031*10^{-3}$ | $1.8032*10^{-3}$ | 136.5 | 0.0168 |
| | | MD | $0.9575*10^{-3}$ | $0.9555*10^{-3}$ | 267.5 | 0.0264 |
| | | RD | $0.5347*10^{-3}$ | $0.5316*10^{-3}$ | 267.5 | 0.0264 |
| | 2 | AD | $1.8027*10^{-3}$ | $1.8029*10^{-3}$ | 142 | 0.0255 |
| | | MD | $0.9665*10^{-3}$ | $0.9617*10^{-3}$ | 267 | 0.0274 |
| | | RD | $0.5484*10^{-3}$ | $0.5412*10^{-3}$ | 267 | 0.0274 |
| | 3 | AFD | 0.1255 | 0.1451 | 151 | 0.0474 |
| 22-232 | 2 | AD | $1.8040*10^{-3}$ | $1.8041*10^{-3}$ | 137 | 0.0175 |
| | | MD | $0.9374*10^{-3}$ | $0.9355*10^{-3}$ | 266 | 0.0294 |
| | | RD | $0.5041*10^{-3}$ | $0.5012*10^{-3}$ | 266 | 0.0294 |
| | 3 | AD | $1.8029*10^{-3}$ | $1.8031*10^{-3}$ | 145 | 0.0316 |
| | | MD | $0.9613*10^{-3}$ | $0.9560*10^{-3}$ | 265 | 0.0316 |
| | | RD | $0.5404*10^{-3}$ | $0.5324*10^{-3}$ | 265 | 0.0316 |

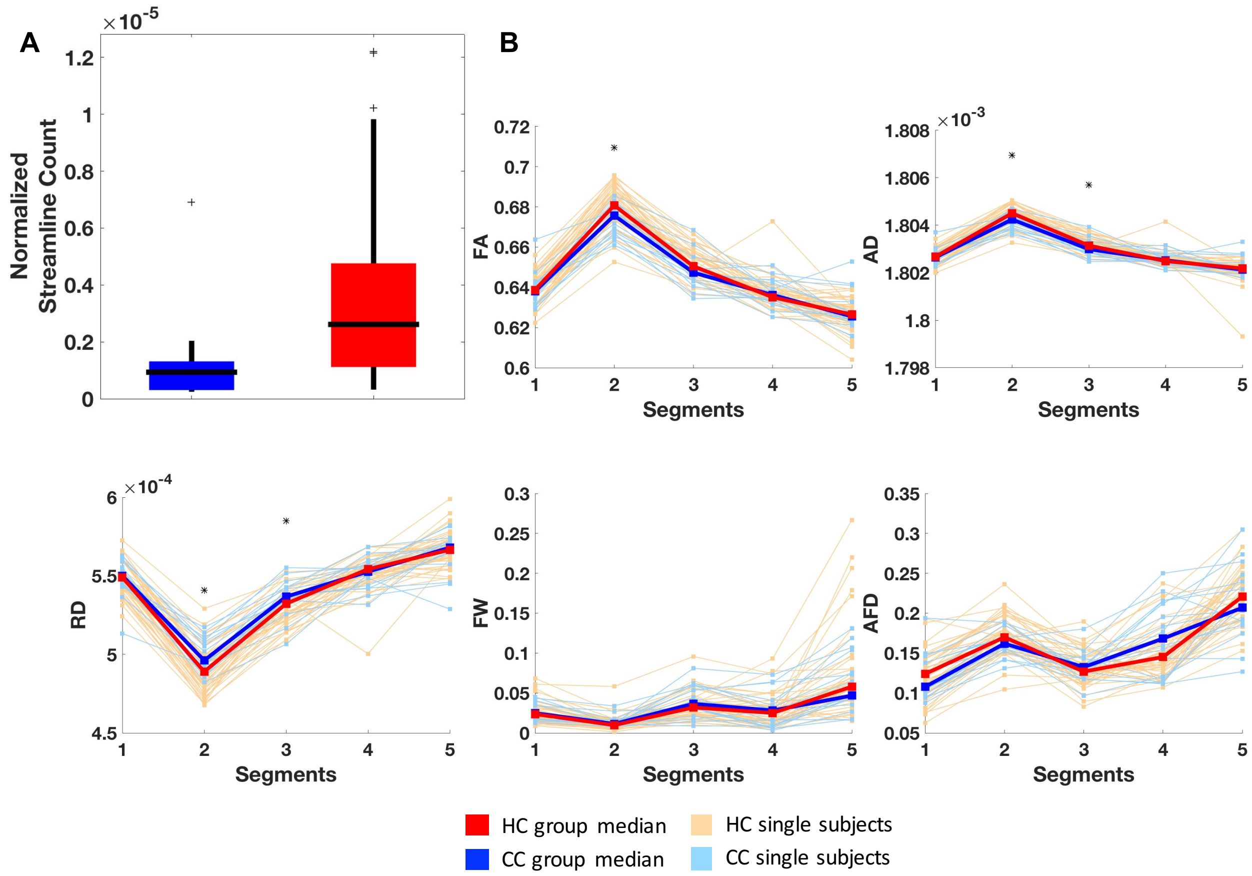


Figure S1. Results of track 22-246. A: Boxplots comparing normalized streamline count between concussed (blue) and healthy control (red) groups. Horizontal black lines illustrate group medians, the edges of the boxes illustrate $75^{th}$ (upper edge) and $25^{th}$ (lower edge) percentiles, vertical black lines illustrate ranges, and black '+' signs illustrate outliers. B: Line plots showing track profiles for AFD, FW, and FW-corrected FA, RD, and AD. Light red and light blue lines indicate median values of each segment for each individual healthy or concussed subject respectively. Dark red and dark blue lines indicate instead group medians at each segment. *: $p<0.05$.

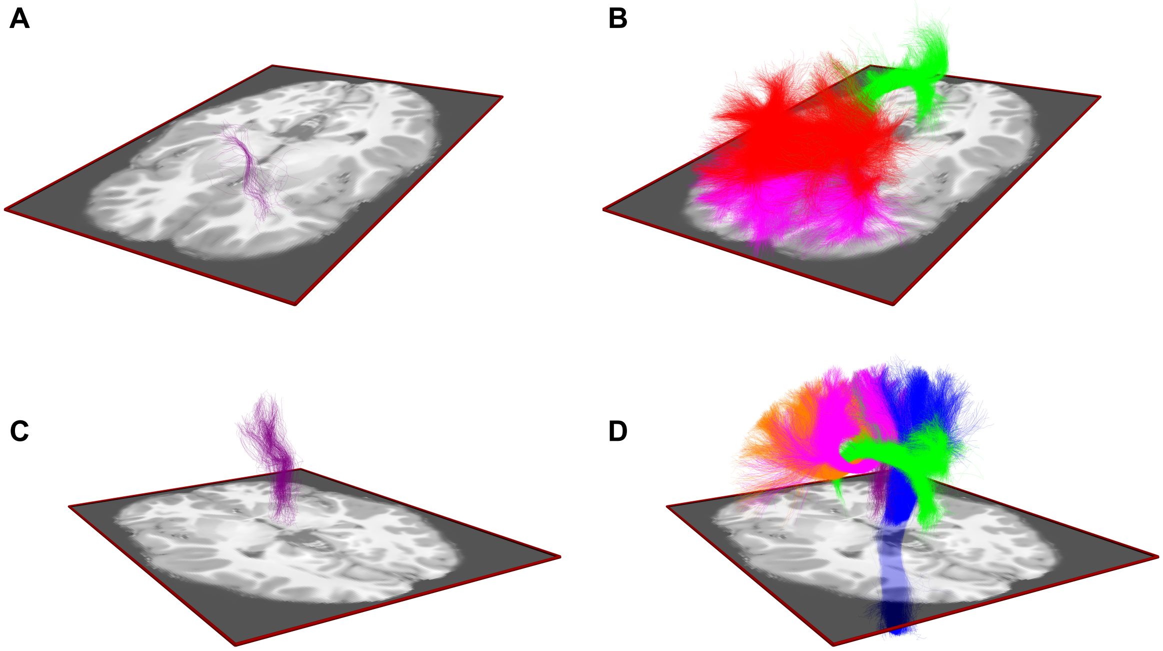


Figure S2. A. Illustration of track 21-179 for one subject, without other tracks visible. B. Illustration of track 21-179 for the same subject, with the cingulum (green) and two segments of the corpus callosum (magenta and red). Track 21-179 is completely occluded by these large tracks. C. Illustration of track 16-246 for one subject, without other tracks visible. D. Illustration of track 16-246 for the same subject, with cingulum (green), corticospinal track (blue), and two other segments of the corpus callosum (magenta and orange). Track 16-246 is still visible.
